# The off-target kinase landscape of clinical PARP inhibitors

**DOI:** 10.1101/520023

**Authors:** Albert A. Antolin, Malaka Ameratunga, Udai Banerji, Paul A. Clarke, Paul Workman, Bissan Al-Lazikani

## Abstract

Four PARP inhibitors have been approved by the FDA as cancer therapeutics, yet there are several clinical settings where no strong rationale exists to inform clinicians on which to choose in terms of clinical effectiveness and toxicity. The four drugs have largely similar PARP family inhibition profiles, but several differences at the molecular and clinical level have been reported that remain poorly understood. We previously hypothesized that PARP inhibitors could have an inherent capacity to inhibit kinases off-target. Here, we characterise the off-target kinase landscape of four FDA-approved PARP drugs. We demonstrate that all four PARP inhibitors have a unique polypharmacological profile across the kinome. Niraparib and rucaparib inhibit DYRK1s, CDK16 and PIM3 at clinically achievable submicromolar concentrations. These represent the most potently inhibited off-targets of PARP inhibitors identified to date and should be investigated further to clarify their potential implications for efficacy and safety in the clinic, including use of PARP inhibitors in combination with other agents, including immunotherapy.

## INTRODUCTION

The demonstration that BRCA1 and BRCA2 mutant human cancer cell lines and tumour xenografts are exquisitely sensitive to small-molecule inhibitors of poly (ADP-ribose) polymerase (PARP) was critical for the clinical development and approval of PARP inhibitors as single agents and provided the first clinical exemplification of synthetic lethality in oncology.^1,2^ All PARP inhibitors currently in the clinic bind to the nicotinamide binding pocket of PARPs through a shared benzamide pharmacophore that is essential for PARP binding, but the individual agents differ in size and flexibility (Figure 1a).^3^ In 2014, olaparib was the first PARP inhibitor to be approved by the FDA for advanced BRCA-mutated ovarian cancer, followed by rucaparib which was licensed for the same indication in 2016 (Table 1).^4,5^ Niraparib was then approved in 2017 as maintenance treatment for recurrent fallopian tube, ovarian and primary peritoneal cancers (Table 1).^6^ In 2018, olaparib and rucaparib also gained approval as maintenance treatment in the same types of cancer while olaparib was additionally licensed for BRCA-mutated HER2-negative breast cancer (Table 1).^4–6^ Most recently, talazoparib was approved for BRCA-mutated HER2-negative breast cancer^7^ (Table 1). Further PARP inhibitors are under clinical development.^1,8,9^ No strong rationale currently exists for selecting one PARP drug over the others in terms of clinical effectiveness and toxicity and prescription is largely based on the approved indication for each drug as well as the reimbursement policy of the relevant healthcare provider.^10,11^ Deeper understanding of the activity and liabilities of individual PARP inhibitors is therefore important to aid clinical decisions and benefit of cancer patients.

**Figure 1.**
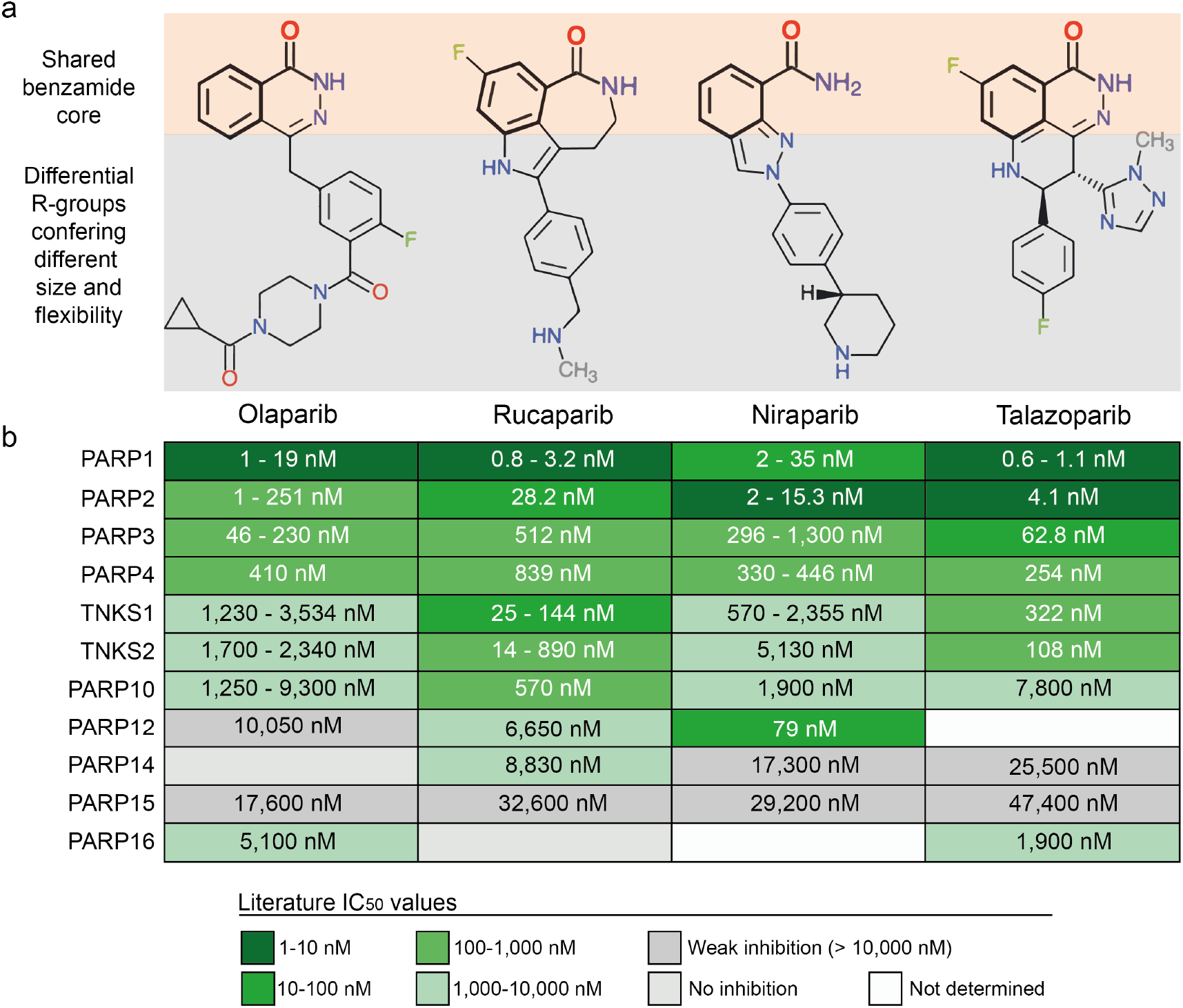
Chemical structures and known PARP activities of clinical PARP inhibitors. **a** Chemical structures of the four FDA-approved PARP drugs. The benzamide core pharmacophore shared by all clinical PARP inhibitors is highlighted in bold and coloured orange. The rest of the chemical structure that is not shared between the inhibitors and confers them with different size and flexibility is coloured grey. **b** Known target profile of clinical PARP inhibitors across members of the PARP enzyme family. IC_50_ values are obtained from the literature and the database ChEMBL and ranges are given where there is more than one published value.^29,35^

**Table 1.**
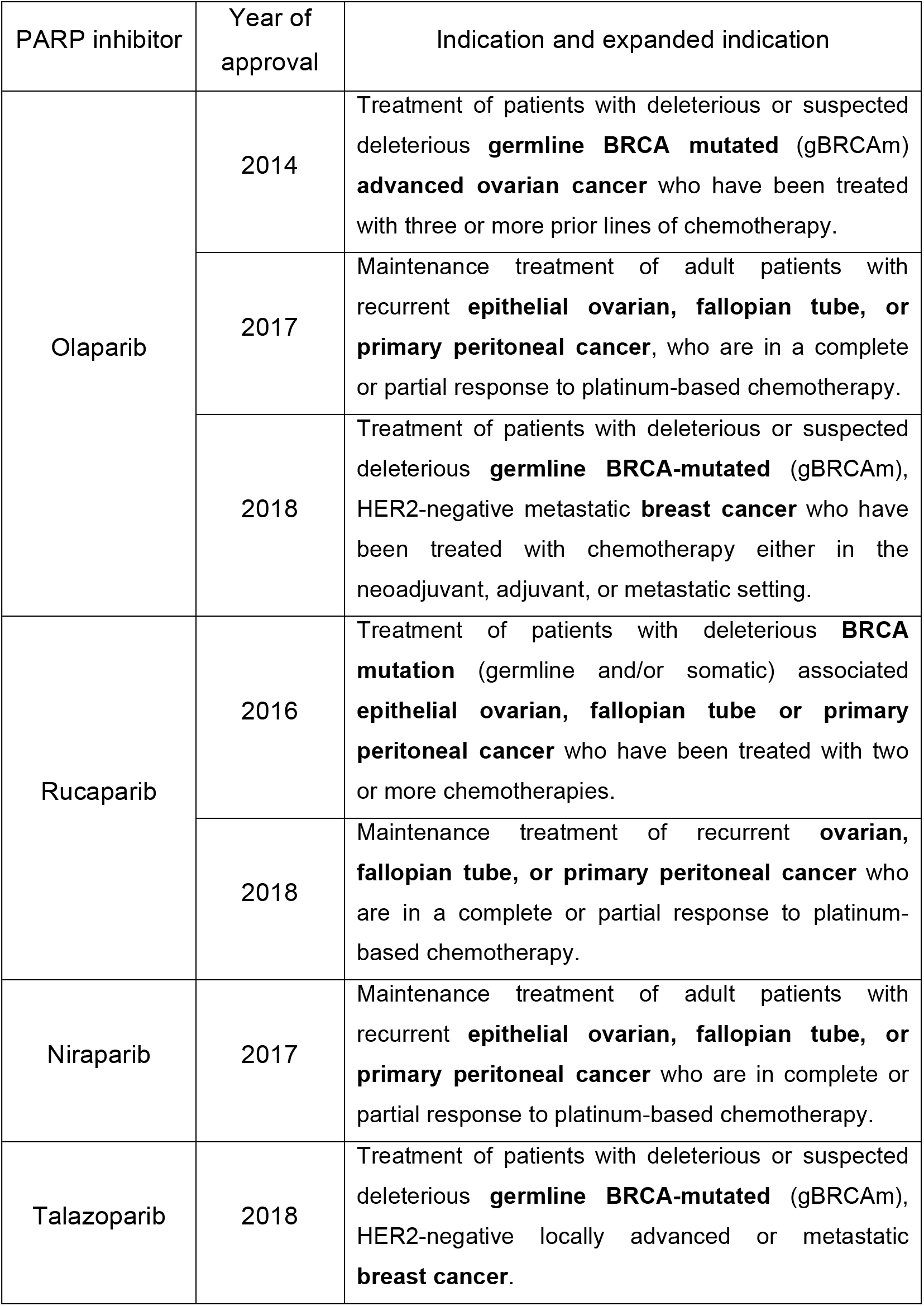
Evolution of the indications for FDA-approved PARP inhibitors. Data extracted from the FDA Hematology/Oncology (Cancer) Approvals & Safety Notifications (accession date 29^th^ May 2018).^65^

| PARP inhibitor | Year of approval | Indication and expanded indication |
| --- | --- | --- |
| Olaparib | 2014 | Treatment of patients with deleterious or suspected deleterious **germline BRCA mutated** (gBRCAm) **advanced ovarian cancer** who have been treated with three or more prior lines of chemotherapy. |
|  | 2017 | Maintenance treatment of adult patients with recurrent **epithelial ovarian, fallopian tube, or primary peritoneal cancer**, who are in a complete or partial response to platinum-based chemotherapy. |
|  | 2018 | Treatment of patients with deleterious or suspected deleterious **germline BRCA-mutated** (gBRCAm), HER2-negative metastatic **breast cancer** who have been treated with chemotherapy either in the neoadjuvant, adjuvant, or metastatic setting. |
| Rucaparib | 2016 | Treatment of patients with deleterious **BRCA mutation** (germline and/or somatic) associated **epithelial ovarian, fallopian tube or primary peritoneal cancer** who have been treated with two or more chemotherapies. |
|  | 2018 | Maintenance treatment of recurrent **ovarian, fallopian tube, or primary peritoneal cancer** who are in a complete or partial response to platinum-based chemotherapy. |
| Niraparib | 2017 | Maintenance treatment of adult patients with recurrent **epithelial ovarian, fallopian tube, or primary peritoneal cancer** who are in complete or partial response to platinum-based chemotherapy. |
| Talazoparib | 2018 | Treatment of patients with deleterious or suspected deleterious **germline BRCA-mutated** (gBRCAm), HER2-negative locally advanced or metastatic **breast cancer**. |

Several differences between individual PARP inhibitors have already been reported at the cellular and clinical levels. When used at micromolar concentrations, differences in DNA strand break repair, phosphorylation of several proteins, cell cycle arrest, and anti-proliferative activities have been described between olaparib, rucaparib and veliparib in cancer cell lines.^12,13^ Moreover, the different capacity of PARP inhibitors to trap PARP at the DNA damage site is widely accepted as important for the mechanism of action of PARP inhibitors but the exact molecular mechanism of this is not completely understood.^14^ Differences in cancer cell sensitivity and synthetic lethality also emerge from recent large-scale profiling experiments, in some instances leading to the prediction of distinct genomic biomarkers (Supplementary Table 1).^15,16^ Overall, it seems important to investigate further the differences between PARP inhibitors and to explore the potential impact on clinical use.

It is now widely accepted that drugs often bind several proteins beyond their intended target ‘polypharmacology’.^17–21^ Moreover, experimental and computational methods continue to uncover previously unknown off-targets of drugs.^19,22–25^ Alongside other factors such as pharmacokinetics, polypharmacology has been demonstrated to cause differences in side-effects between drugs of the same class.^26^ In addition, understanding of polypharmacology can lead to the exploitation of drugs in novel indications, such as the recent approval of crizotinib in ROS1-driven non small-cell lung cancer.^26–28^

The selectivity and polypharmacology of PARP inhibitors within the PARP-family was recently characterised *in vitro* using an enzymatic inhibition assay.^29^ Veliparib and niraparib were shown to be more selective for PARP1 and PARP2 compared to olaparib, rucaparib and talazoparib which show broader pan-PARP activity (Figure 1b).^29^ However, this differential intra-family PARP selectivity is insufficient to explain all the differences observed between clinical PARP inhibitors. In 2014, we reported for the first time that the different polypharmacology patterns between PARP inhibitors extended beyond the PARP enzyme family.^30^ We demonstrated that rucaparib inhibited at least nine kinases with micromolar affinity whereas veliparib inhibited only two kinases and olaparib did not exhibit activity against any of the 16 kinases tested.^30^ During a high throughput screen for RPS6KB1 kinase inhibitors we identified a series of carboxamidobenzimidazoles that were confirmed to bind RPS6KB1 by orthogonal methods including X-ray crystallography.^31^ The carboxamidobenzimidazoles are known inhibitors of PARP^32^ and the existence of a crystal structure of a carboxamidobenzimidazole bound to RPS6KB1 kinase^31^ prompted our speculation that all PARP inhibitors could have an intrinsic capacity to inhibit kinases. This activity could result from the ability of their shared benzamide pharmacophore to interact with the highly-conserved kinase hinge region (Figure 1a).^30,31^ This suggested that – depending on its individual molecular size and decoration – each PARP inhibitor could have a unique off-target kinase profile that may remain as yet unexplored and would be important to characterise.^21,30^ More recently, an unbiased, large scale, mass spectrometry-based chemical proteomics approach uncovered new, low-potency affinities of the PARP inhibitor niraparib.^33^ However, the chemical proteomics approach used was not able to reproduce published, stronger off-target kinase interactions.^30^ This illustrates the limitations of any single method for comprehensively identifying drug polypharmacology.^30^

Here, we use computational and experimental methods to characterise the off-target kinase landscape of the four FDA-approved PARP inhibitors, olaparib, rucaparib, niraparib and talazoparib. We uncover novel, submicromolar off-target kinases – which to our knowledge represent the most potent off-target interactions of PARP inhibitors identified to date. We have also compiled and review the clinical side-effect profiles of the approved PARP inhibitors. Our findings further demonstrate the unique kinase polypharmacology of each PARP inhibitor that investigators should be aware of, and which could potentially open new avenues for the differential exploitation of clinical PARP inhibitors in precision medicine.

## RESULTS

### In silico target profiling predicts new kinase off-targets of clinical PARP inhibitors

We applied three parallel computational methods to predict off-targets: 1) a consensus of six ligand-based chemoinformatic methods integrated in the Chemotargets CLARITY platform^34^; 2) the Similarity Ensemble Approach (SEA)^26^; and 3) the multinomial Naive Bayesian multi-category scikit-learn method implemented in ChEMBL.^35^ The common principle for these methods is that chemically similar molecules should share similar bioactivity profiles against molecular targets; however, the details of the methods, including the representation of compounds and similarity calculations used, are distinct. We used these three methods to predict the kinase off-targets of the four FDA-approved PARP inhibitors, olaparib, rucaparib, niraparib and talazoparib. In addition to recovering most of the known interactions with members of the PARP family, the three methods predicted a total of 58 interactions between PARP inhibitors and kinases, with only 10 of them being previously known^30^ (Table 2, Supplementary Tables 2-4).

**Table 2.**
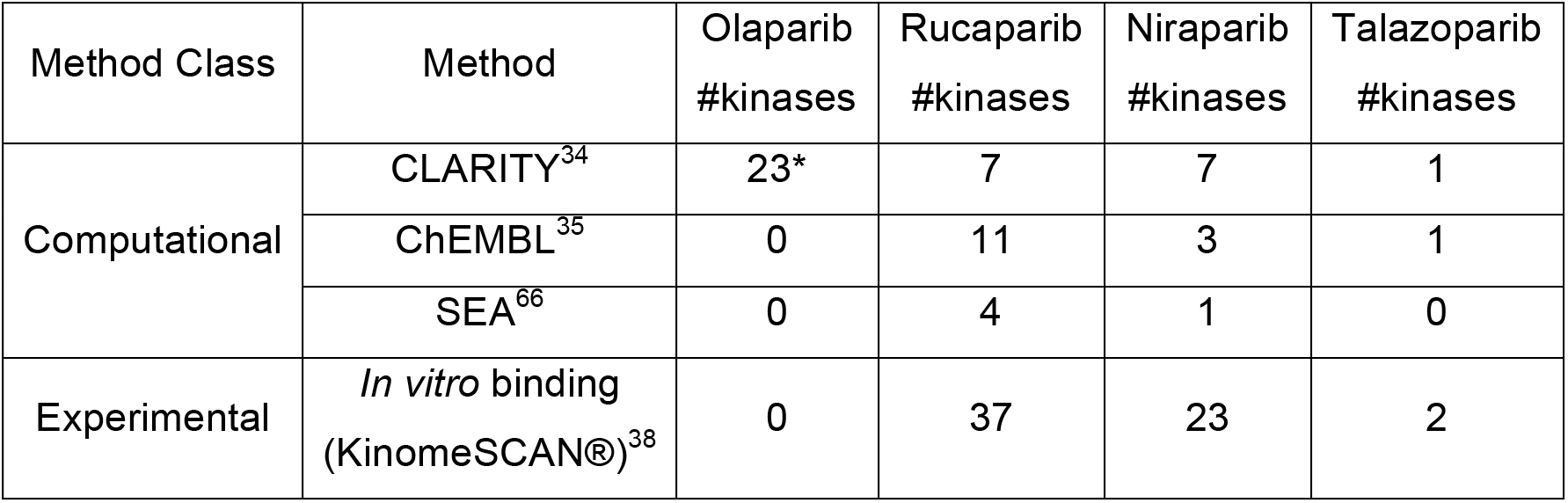
Comparison between the number of kinases predicted for clinical PARP inhibitors by three *in silico* target profiling methods and experimentally identified by *in vitro* kinome binding at 10 μM. *Prediction originating from the similarity to a single kinase inhibitor that is likely a false positive.

CLARITY predicted 23 kinases as potential off-targets of olaparib (Supplementary Table 2). However, neither ChEMBL nor SEA predicted any kinase for this PARP drug (Supplementary Tables 3-4). A close inspection of the CLARITY predictions revealed that they were all generated from the similarity of olaparib to a single kinase inhibitor that was likely to be a false positive due to the absence within its structure of a benzamide moiety, which is known to be important for PARP binding (Supplementary Table 2).^3^

CLARITY predicted seven kinases as potential off-targets of niraparib while ChEMBL predicted three kinases and the SEA method predicted one kinase (Table 2, Supplementary Tables 2-4). However, while all the methods predicted kinases as potential off-targets for niraparib, no two methods predicted the same kinase. These predictions might reflect that niraparib has general kinase-binding features that are not specific to any single kinase, since there is no agreement between the methods on the specific kinases. However, it is still interesting that all the three methods predict kinases as potential off-targets of niraparib, particularly given that the only known kinase off-target of niraparib reported previously in the literature had low affinity (DCK IC_50_ = 67.9 μM).^33^ In addition, some of the kinase inhibitors that are identified as similar to niraparib present PARP-binding features. For example, CHEMBL2035040, a weak AKT inhibitor, shares the key benzamide moiety with niraparib and other PARP inhibitors.

We have previously demonstrated that rucaparib inhibits nine kinases with micromolar potency.^30^ Given the high degree of polypharmacology of many kinase inhibitors, we hypothesized that rucaparib could inhibit more kinases than the ones already identified. None of the three computational methods used here predicted the nine kinase off-targets of rucaparib that are already known. CLARITY, ChEMBL and SEA correctly predicted four, two and four known kinase off-targets, respectively, for rucaparib. Additionally, CLARITY predicted three new kinases, and ChEMBL predicted nine new kinase off-targets (Table 2, Supplementary Tables 2-4). Overall, twelve new kinase off-targets in total were predicted for rucaparib.

Finally, CLARITY and ChEMBL predicted only one kinase off-target each for talazoparib whilst SEA predicted none (Table 2). This low number of kinase off-target predictions suggests that it less likely that talazoparib inhibits kinases (Supplementary Tables 2-4).

Overall, the lack of consensus on specific kinase off-targets between the three computational methods (Table 2, Supplementary Tables 2-4) is noteworthy and requires future exploration of the methodology, especially given the expanding use of these approaches.^36^ It is particularly puzzling given that all three methods are based on chemical structure similarity and use the same underlying medicinal chemistry databases – highlighting the sensitivity of such predictions to the specific computational representation of compounds and statistics utilised. Based on our experience, we advise that, until these methodologies improve, researchers should apply as many predictive computational methods as possible. However, despite the differences in the detail, an important message is that all three methods did predict a range of kinases as potential off-targets of niraparib and rucaparib, thus increasing the confidence in our hypothesis that additional kinase off-targets are likely to be found for PARP drugs (Table 2).

### Kinome profiling with a binding assay uncovers differential polypharmacology between clinical PARP inhibitors

To follow up the computational analysis, we performed a comprehensive *in vitro* kinome screen. To do this we employed the DiscoveRx (https://www.discoverx.com) KinomeScan *in vitro* binding technology^37,38^ that has been widely used for kinome profiling in drug discovery. At the time of conducting our screen, this kinome panel was the largest commercially available and comprised 468 *in vitro* binding assays corresponding to 392 unique human kinases (76% of the human kinome)^39^ (Supplementary Table 5). Also included were assays comprising mutated, deleted, phosphorylated or autoinhibited forms of proteins (n = 63), secondary or pseudokinase domains (n = 8), non-human isoforms (n = 3) and CDK complexes with different cyclins (n = 2) (Supplementary Table 5). The assays were performed at a single relatively high concentration of 10 μM to identify initially both low and high potency off-targets.

The results of the use of the *in vitro* binding assay for kinome profiling reveal marked differences between the kinase polypharmacology of PARP inhibitors. As illustrated in Figure 2, rucaparib and niraparib bind to many kinases while talazoparib binds only to two kinases with modest affinity and olaparib does not bind to any of the 392 kinases tested (Supplementary Table 5). The technology used involves a competition assay, defining activity as ≥65% of the kinase being competed off an immobilised ligand at 10 μM^38^. Using this measure, rucaparib binds to 37 kinases while niraparib binds to 23 kinases (Table 2, Supplementary Table 5). Moreover, there is only partial overlap between the measured kinase polypharmacology of niraparib and rucaparib, with both drugs binding to 15 shared kinases. Interestingly, the two kinases to which talazoparib showed weak binding are CLK3 and MTOR, neither of which bind the more broadly acting rucaparib and niraparib.

**Figure 2.**
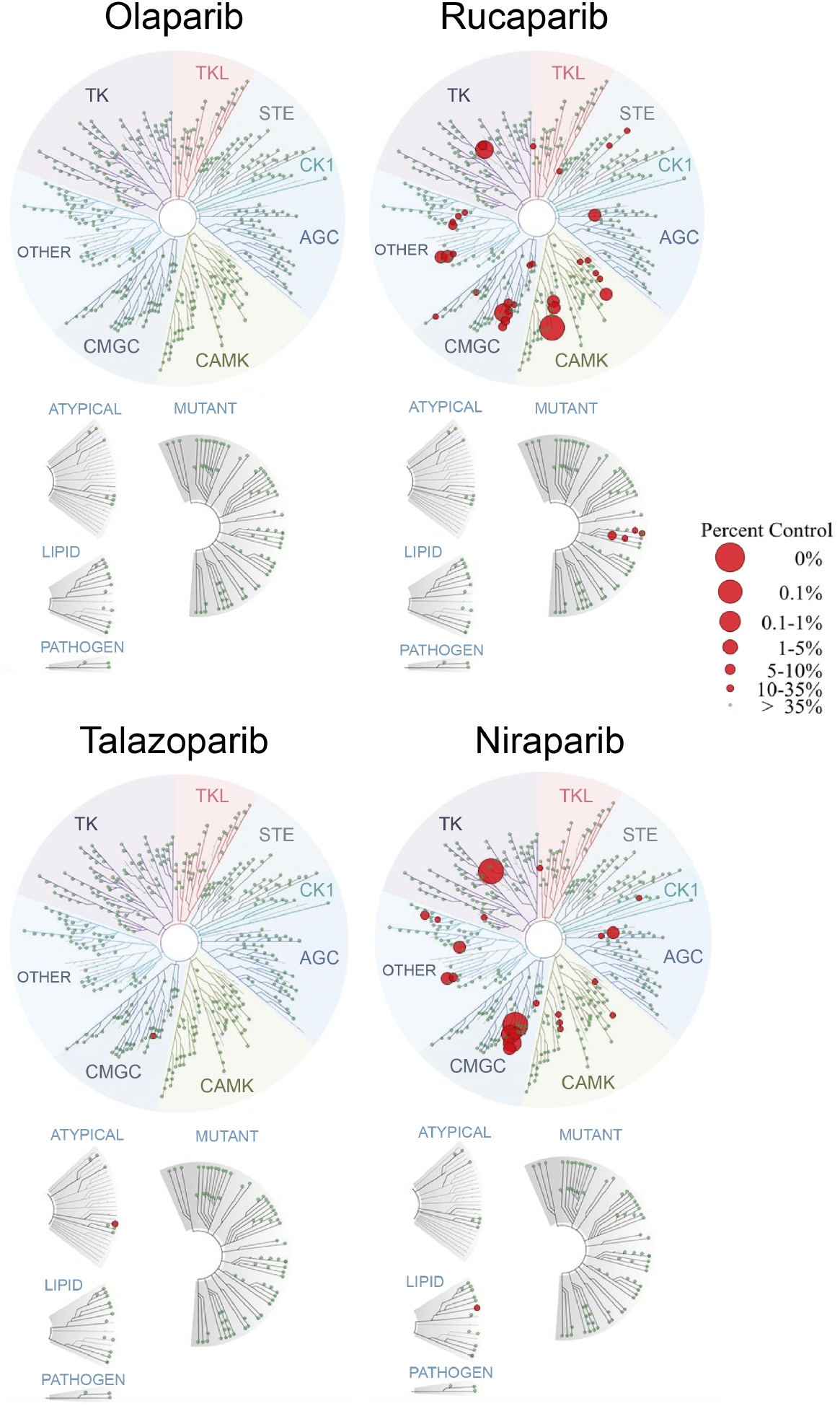
Kinome profiling of the four FDA-approved PARP inhibitors across 392 unique human kinases and 76 mutated, atypical and other forms. This was carried out using the *in vitro* binding platform of DiscoveRx’s KinomeScan®.^38^ The assays were perfomed at a single 10 μM concentration. The TREEspot™^64^ representations of the kinome tree, with superimposed *in vitro* binding data for each PARP inhibitor, illustrate how rucaparib and niraparib bind to a significant number of kinases while talazoparib only modestly binds to two kinases and olaparib does not bind to any of the kinases tested.

Importantly, as illustrated in Figure 2, the off-target kinase activities of the four PARP inhibitors do not cluster in one single kinase family but are fairly widely distributed across the kinome. When carried out at the 10 μM concentration used, the *in vitro* binding assay is able to recover most previously known off-targets of rucaparib, including the most potent known interactions with PIM1 and DYRK1A. However, the previously described weaker interactions of rucaparib with PRKD2, CDK9, PIM2 and ALK were not reproduced by the binding assay. A further two known off-targets, CDK1 and DCK, were not available in the kinome panel used.

Overall, our results provide empirical evidence that the polypharmacology profile is indeed different for the four different PARP inhibitors studied. Consistent with the prediction by the computational methods, rucaparib and niraparib demonstrate multiple kinase polypharmacology, with less or no off-targets for talazoparib and olaparib.

### Orthogonal catalytic inhibition assay confirms DYRK1s, PIM3 and CDK16 as submicromolar off-targets of niraparib and rucaparib

Next, we validated the observed activities in an orthogonal experimental method, namely direct inhibition of kinase catalytic activity using Reaction Biology’s HotSpot platform (http://www.reactionbiology.com).^40^ This platform employs a widely-used and validated radiometric assay which measures inhibition of catalytic activity.^41^ Of the 24 kinases showing the greatest binding by the PARP inhibitors (≥85% binding at 10 μM, Supplementary Table 5) four were not available for follow-up testing at Reaction Biology (Highlighted in Supplementary Table 5). Thus, in total we tested 20 kinases in a radiometric catalytic inhibition assay, initially at a 1 μM of the PARP inhibitors (Table 3). Of these, four were inhibited more than 50% by rucaparib and/or niraparib, namely DYRK1B, CDK16/cyclin Y, PIM3 and DYRK1A (Table 3).

**Table 3.** Validation of the kinase binding hits with greater binding effect at 10 μM from the kinome profiling (Figure 2, Supplementary Table 5) using an orthogonal radiometric catalytic assay. The assay directly measures kinase catalytic activity using a widely-validated radiometric assay.^40^ The table displays the average of duplicate (n = 2) measurements of the percentage of enzyme inhibition relative to DMSO controls sorted by maximum percentage of inhibition. All assays were performed using 1 μM of drug concentration and Km concentration of ATP. From all the tested kinases, only 4 inhibit the enzyme by >50%. These most potent interactions, expected to be submicromolar, are displayed in bold. n.b. not binding (Supplementary Table 5). n.d. not determined due to low binding (Supplementary Table 5). * n = 1.

| Kinase Group | Gene Name | Kinase complex and aliases | Niraparib | Rucaparib |
| --- | --- | --- | --- | --- |
| **CMGC** | **DYRK1B** | - | **82%** | **53%** |
| **CMGC** | **CDK16** | **CDK16/cyclin Y (PCTAIRE, PCTK1)** | n.b. | **82%** |
| **CAMK** | **PIM3** | - | 15% | **78%** |
| **CMGC** | **DYRK1A** | **DYRK1** | **76%** | 49% |
| CMGC | HIPK1 | - | 40% | -16% |
| CAMK | MYLK4 | - | 19% | 36% |
| Other | AURKB | Aurora B | 34% | n.b. |
| CMGC | HIPK2 | - | 23% | -6% |
| CAMK | PIM1 | - | 22% | 27%* |
| CMGC | CSNK2A1 | CK2a | 12% | 21% |
| AGC | LATS2 | - | 17% | n.b. |
| CMGC | CSNK2A2 | CK2a2 | -1% | 17% |
| AGC | CIT | STK21 | 10% | 15% |
| Other | HASPIN | Haspin | 10% | -6% |
| CMGC | CDK4 | CDK4/cyclin D3 | n.b. | 10% |
| CMGC | HIPK3 | - | 5% | -12% |
| CAMK | TSSK3 | STK22C | n.b. | -1% |
| PKL | PIK3C3 | VPS34 | -1%* | n.b. |
| CAMK | PIM2 | - | -5% | n.b. |
| CAMK | STK17A | DRAK1 | n.d. | -6% |

Some of the initial hits identified in the DiscoveRx KinomeScan binding assay – such as TSSK3 – could not be reproduced using the orthogonal radiometric catalytic assay. This is frequently observed when comparing binding and catalytic assays,^42^ due to factors such as differences in assay methods and conditions (e.g. protein constructs and drug concentrations used). Interestingly, although the primary hits from the binding assay are widely distributed across the kinome tree (Figure 2), the 20 selected kinases showing the greatest binding by the PARP inhibitors are from only 5 different kinase groups and the submicromolar kinase off-targets of PARP inhibitors in the radiometric catalytic assay are all from the CMGC and CAMK groups (Table 3). At the submicromolar level tested in the catalytic assay (> 50% inhibition at 1 μM), rucaparib inhibited kinases from both groups but niraparib inhibited only two kinases from the CMGC group. DYRK1B was the only kinase inhibited by both rucaparib and niraparib at concentrations below 1 μM (Table 3). These results further emphasize the different kinase polypharmacology behaviour exhibited between these two clinical PARP inhibitors.

Given our identification of potent new submicromolar off-targets that could have clinical implications, we measured the IC_50_ for the submicromolar kinases in the catalytic assay (> 50% inhibition at 1 μM) using Reaction Biology’s radiometric 10-point concentration-response inhibition assay (Figure 3). Rucaparib inhibits three kinases (CDK16, PIM3 and DYRK1B) with submicromolar IC_50_ values, the most potent being CDK16 (IC_50_ = 381 nM). In contrast, niraparib inhibits only two kinases with submicromolar IC_50_ values, the most potent being DYRK1B (IC_50_ = 254 nM). To our knowledge, this is the first report of submicromolar non-PARP family off-targets of PARP inhibitors.

**Figure 3.**
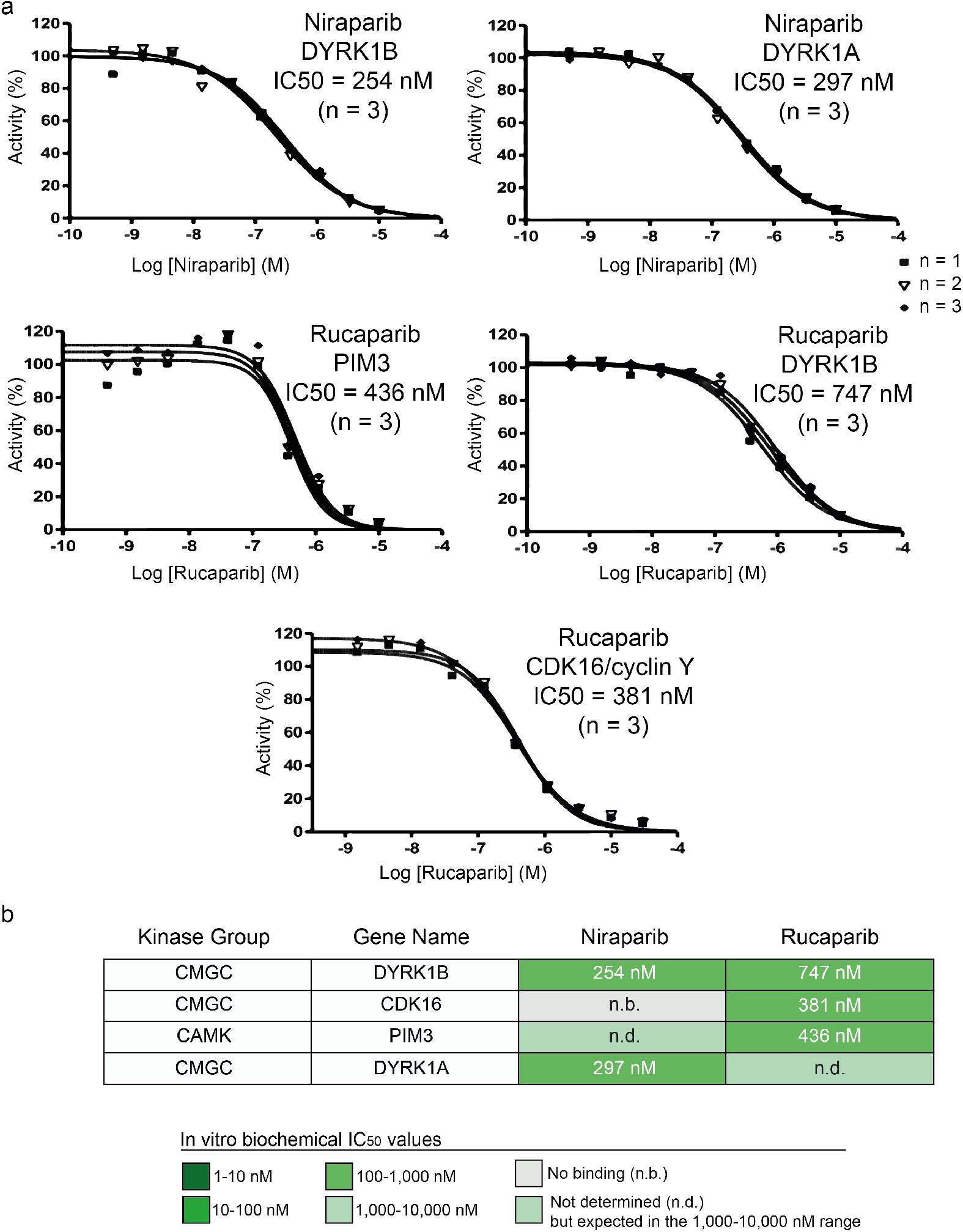
Concentration-response curves and IC_50_ calculation for the most potent kinase off-target interactions of clinical PARP inhibitors. **a** concentration-response curves of the interactions with greater binding effect at 10 μM of niraparib (top) and rucaparib (bottom) analysed in triplicate using Reaction Biology’s HotSpot radiometric assay that directly measures kinase catalytic activity.^40^ **b** Table summarising the calculated IC_50_ values for the kinase off-targets DYRK1B, CDK16, PIM3 and DYRK1A.

### Docking experiments to study PARP-kinase polypharmacology at the atomic level

To attempt to understand the underlying basis of PARP-kinase polypharmacology at the atomic level we used *in silico* docking to predict the binding modes of clinical PARP inhibitors for their most potent kinase off-targets.

Of the four protein kinases inhibited with submicromolar affinities by rucaparib or niraparib (Figure 3), only DYRK1A and CDK16 had a 3D structure deposited in the PDB.^43^ Using GOLD (https://www.ccdc.cam.ac.uk/solutions/csd-discovery/components/gold/), we compared the binding poses of all four clinical PARP inhibitors against these two kinases (Figure 4 and Supplementary Figure 1). Docking scores using GOLD’s S(PLC) scoring function are concordant with kinase binding affinity (Figure 5 and Supplementary Figure 1). In order to facilitate the analysis of the molecular interactions between PARP inhibitors and kinase proteins in the best GOLD scoring poses, we used LigPlot+ to generate 2D schematic diagrams of these protein-ligand interactions.

**Figure 4.**
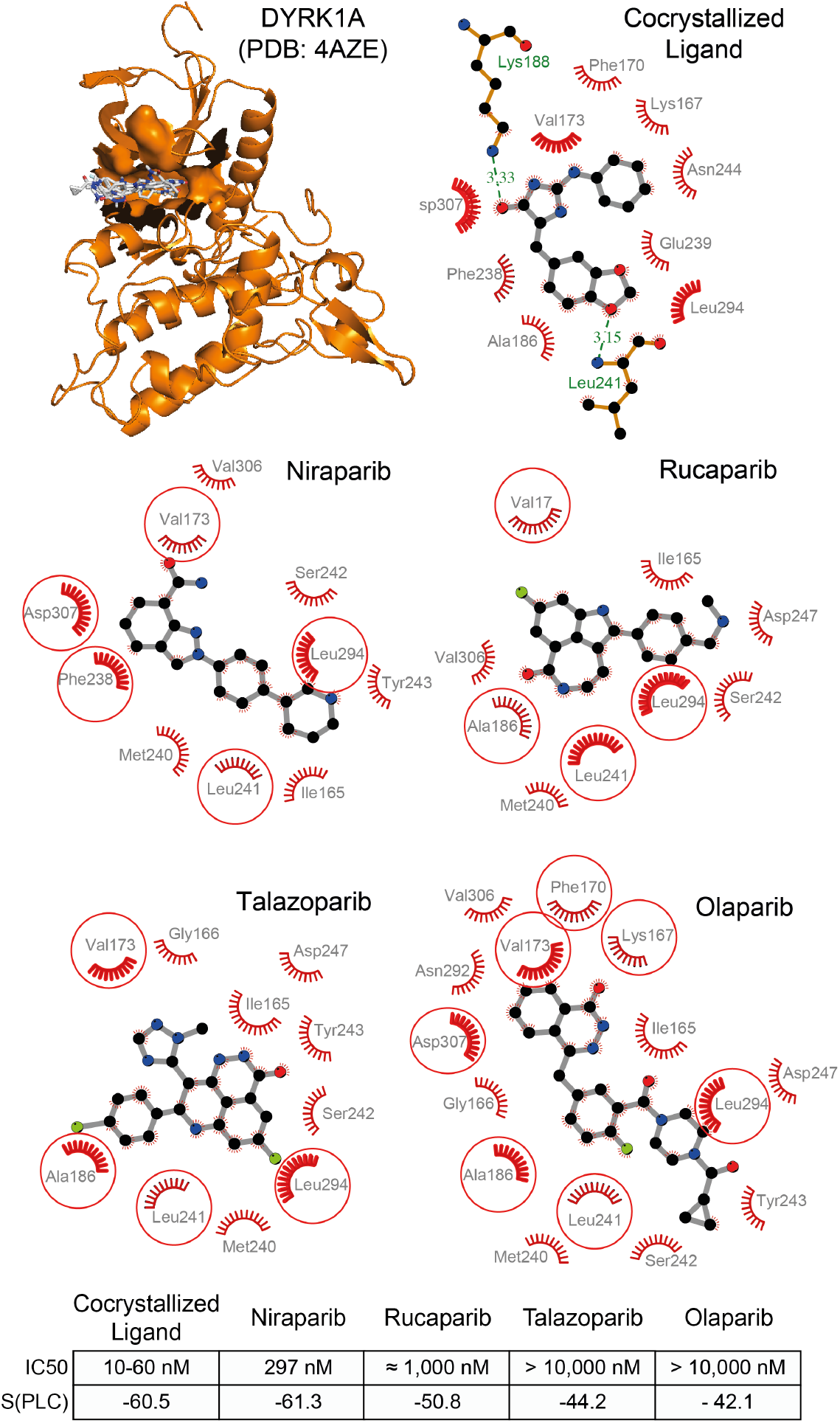
Docking poses with the top GOLD S(PLC) score of four clinical PARP inhibitors in DYRK1A kinase. In the top left panel, all ligands are superimposed and the whole protein structure is displayed using MacPyMOL (PyMOL v1.8.0.6). In subsequent panels, LigPlot+^44^ was used to generate schematic diagrams of protein-ligand interactions for the cocrystalized ligand and clinical PARP inhibitors. The lower Table summarises the value of the GOLD scoring function for the highest-scoring pose – the one represented – and the IC_50_ determined in this work (Figure 3) or extracted from the literature for each of the ligands.

**Figure 5.**
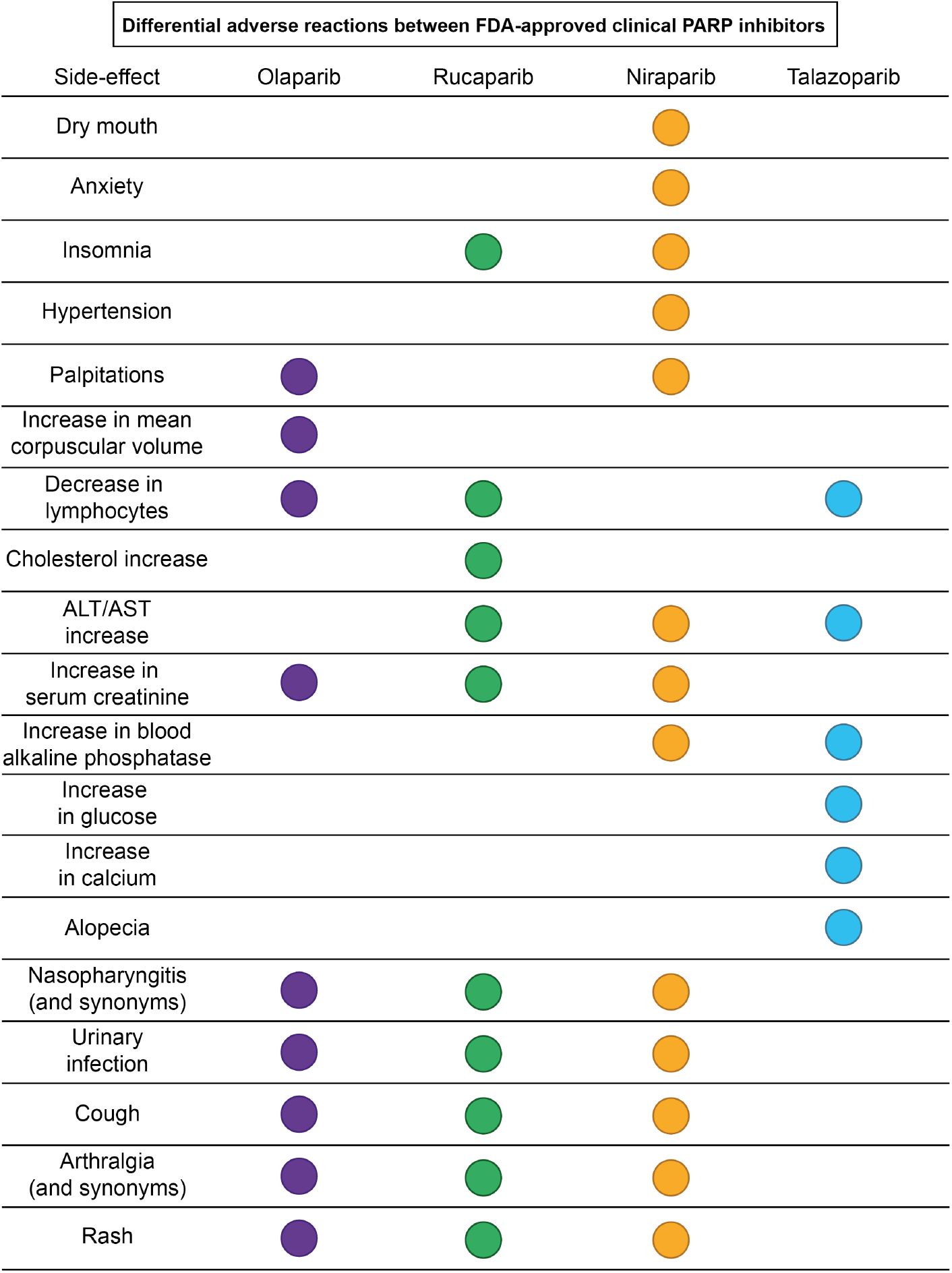
The nineteen differential adverse reactions between FDA-approved clinical PARP inhibitors extracted from the FDA prescribing information and published results of the largest clinical trials. Side-effect frequencies are not considered due to the differences between the cut-offs used in each trial and FDA prescribing information for each PARP inhibitor (see Methods for details). Each adverse reaction considered common for at least one PARP drug and not identified for at least another PARP inhibitor is represented as a circle. The circles are coloured according to the drugs that present this adverse reaction in their prescribing information or publication of their largest clinical trial.

Additionally, we analysed the structures of the two kinases co-crystalized with inhibitors that share the cyclic benzamide moiety found in most PARP inhibitors (Supplementary Table 7).^44^ In the case of DYRK1A, no hydrogen-bond interaction was formed in the best scoring GOLD pose of the PARP inhibitors olaparib, rucaparib, niraparib and talazoparib with the kinase. In contrast, the cocrystalized inhibitor forms two hydrogen bonds with DYRK1A (Figure 4). In the case of CDK16, only rucaparib and talazoparib form one hydrogen bond each while the cocrystalyzed ligand interacted via four hydrogen bonds with CDK16 (Supplementary Figure 1).

There was no evident relationship between the number of overall interactions in the best docking poses and the affinity of each PARP inhibitor for CDK16 or DYRK1A. The lack of shared contacts demonstrates that it is unlikely that the benzamide moiety, which is a key feature for PARP binding, is a major contributor to kinase binding.^30^ This is consistent with our experimental results as all our tested PARP inhibitors contain this moiety. It is possible that the flatter shape of niraparib and rucaparib might allow a better fit in the kinase binding sites as compared to the bulkier talazoparib and the more flexible olaparib. In turn, this might be responsible for their better docking scores and affinities for kinases. However, experimental validation of this binding is required to confirm any of these hypotheses.

### Analysis of clinical data shows differential side-effects between PARP inhibitors

As shown in Figure 1, all examined PARP inhibitors have low nanomolar potencies against their PARP targets whereas the potencies against kinases are in the 250 nM to low micromolar range. Moreover, drug penetration inside solid tumours is frequently limited.^45^ Therefore, at the potencies identified here, it is unlikely that the inhibition of kinases by rucaparib and niraparib contributes to therapeutic activity at the tumour site.

However, we reasoned that submicromolar kinase off-targets activities are sufficiently potent to have potential additional, clinically relevant pharmacological effects in better perfused organs outside the immediate tumour site. Rucaparib and niraparib received FDA-approval in 2016 and 2017 for a recommended dose of 600mg taken twice daily and 300mg taken once daily, respectively (Table 1).^5,6^ At these clinical doses, their steady-state Cmax concentrations in plasma range between 2-9 μM for rucaparib^46^ and 3-4 μM for niraparib.^47^ These micromolar Cmax concentrations are well above the submicromolar *in vitro* IC_50_ concentrations for their most potent kinase off-targets.

There are no clinical trials comparing PARP inhibitors directly side by side. Consequently, we first used the FDA prescribing information^4–6^ to analyse all 61 reported adverse events and laboratory abnormalities for the four FDA-approved PARP inhibitors (Supplementary Table 8). Given the lack of direct, quantitative, comparative studies, we summarised the findings into qualitative categories to allow comparison (Supplementary Table 9). Next, we abstracted the reported adverse reactions in the four largest clinical trials of olaparib,^48^ rucaparib^49^, niraparib^50^ and talazoparib^51^. Despite obvious limitations in comparing different trials, the combination of the abstracted information provides high-level insights into similarities and differences between the different PARP inhibitor side-effects (detailed in Supplementary Tables 8-9).

Of the 61 analysed parameters, 21 are shared between all four approved PARP inhibitors although some are only rarely observed. These include frequently observed side-effects of cancer therapeutics such as nausea, vomiting or diarrhoea. Of the 61 parameters, 40 were reported to be commonly observed side-effects for at least one of the four drugs (Supplementary Table 9). Several of these effects are shared between most PARP drugs, such as the increase in serum creatinine, which is reported during treatment with olaparib, rucaparib and niraparib but not talazoparib (Figure 5). Other side-effects, such as palpitations, are reported only for two clinical PARP inhibitors. Finally, all four drugs have unique commonly observed side-effects not shared with other PARP inhibitors. For example, rucaparib is the only PARP inhibitor reported to increase cholesterol and talazoparib is the only PARP inhibitor reported to produce alopecia (Figure 5). All these nineteen distinct side-effects are commonly observed for their respective drugs (as reported in the Prescribing Information). Overall, FDA-approved PARP drugs appear to have distinct side-effect profiles and we hypothesize that their unique polypharmacological profiles could contribute to them.

## DISCUSSION

In this study, we computationally predict and experimentally characterise the off-target kinase landscape of the four clinical PARP inhibitors that are currently approved. We demonstrate that each PARP inhibitor has a unique off-target profile across the kinome. We uncover novel kinase off-targets for the FDA-approved PARP drugs niraparib and rucaparib which we experimentally confirm to have submicromolar inhibitory activities (Figures 2 and 3). Niraparib inhibits DYRK1A and DYRK1B whilst rucaparib inhibits CDK16, PIM3 and DYRK1B – all with submicromolar affinities (Figure 3). We propose that the inhibition by niraparib and rucaparib of DYRK1A, CDK16 and PIM3, among other kinases, may have potential clinical relevance and thus warrant further investigation. Moreover, our findings highlight the importance of considering kinase off-targets in the future discovery and development of PARP inhibitors.

Our results also illustrate the challenge of comprehensively uncovering drug polypharmacology. We find limitations in all screening assay formats, including differences seen with computational prediction methodologies and also chemical proteomics. For example, although all computational methods that we used predicted kinases as potential off-targets of PARP inhibitors, the methods showed little overlap either in the precise computational predictions or with the results of our experimental data (Supplementary Tables 2-6). It is important to note that currently available bioactivity data in public databases are strongly biased towards commonly studied targets, including many kinases. This bias may well contribute to the strong computational prediction of kinase polypharmacology that we observe in this study. Increasing the target coverage of the public databases will improve the training of computational models. Meanwhile, users of such methods are advised to apply as many methods as possible in order to seek consistent predictions and to rely only on those that are experimentally validated.

The availability of experimental *in vitro* target profiling panels at contract research organisations is strongly enabling – especially democratising off-target identification and validation for smaller enterprises and academic groups. However, technological differences between platforms, including expression constructs and systems, purification procedures and assay conditions, affect the results found.^42^ In this study, most of the already known off-targets of PARP inhibitors were reproduced by the *in vitro* competitive binding platform that we used for initial profiling. However, in line with previous observations,^42^ some of the strongest binding signals at 10 μM were not reproduced using a well-established radiometric catalytic assay. Moreover, although increasing in coverage, current screening panels are not yet fully comprehensive – even across widely-studied target families such as kinases.^52^ Finally, the results of a recent chemical proteomics analysis of the selectivity of clinical PARP inhibitors failed to identify our experimentally confirmed findings – but of course are able to sample a broader fraction of the proteome.^33^ There are several factors limiting the use of chemical proteomics in this setting, such as the level of expression of proteins in the cells used and the unknown full effects of the attached tags across the proteome.

It is clear that no method is free of limitations. Overall, our results illustrate the complementarity between different methods in addressing the challenging task of systematically uncovering the molecular target profile of drugs to further our understanding of polypharmacology and its potential impact for efficacy and safety in the clinic.

The clinical PARP inhibitors exhibit different sensitivity across cancer cell lines when measured in large-scale screens, enabling the prediction of distinct genomic biomarkers of drug sensitivity (Supplementary Table 1).^53^ It is possible that differential effects between cell lines may relate to the different polypharmacology of PARP inhibitors. Both the observed differential cellular effects and the differential polypharmacology necessitate consideration of more system-wide effects of PARP inhibitors and their kinase off-targets, particularly in organs that are exposed to high drug concentrations, such as the blood and the liver. Indeed both DYRK1A and CDK16 proteins are highly expressed in the bone marrow, immune cells and the liver while CDK16 is very broadly expressed.^54^

As discussed above, rucaparib and niraparib display a high degree of selectivity for PARPs (low nanomolar) over kinases (high nanomolar). Therefore, at the tumour site *in vivo*, these off-target activities are unlikely to have significant anti-tumour effect. However, the clinical concentrations achieved, especially at sites of higher drug concentrations including the liver, may have potential clinical implications for the off-target activities.

We analysed the FDA prescribing information and data from major clinical trials for the four FDA-approved PARP inhibitors and identify differences in their side-effects. The comparison shows that each PARP drug has a unique clinical side-effect profile and we hypothesize that this may potentially relate to the unique off-target profile (Figure 5, Supplementary Tables 8-9). Since side-effects could result through a multitude of mechanisms, it is not possible to ascribe each side-effect to specific off-targets. Nevertheless, it is tempting to speculate that the unique inhibition of PIM3 by rucaparib and not by olaparib, niraparib or talazoparib may contribute to the unique elevations in cholesterol that are observed in patients treated with rucaparib but not other three PARP inhibitors (Figure 3). The only PARP family member for which rucaparib shows greater affinity than the rest of the PARP inhibitors, and which has been associated with cholesterol homeostasis, is PARP10.^55^ However, the small difference in affinity for PARP10 (Figure 1) between olaparib, niraparib and talazoparib (below 4-fold in the case of niraparib) does not strongly support this hypothesis. Interestingly, PIM3 has been recently found to be regulated downstream of mTORC1 by miR-33 – encoded by the SREBP loci.^56^ SREBP and miR-33 are known regulators of cholesterol homeostasis.^57^ Moreover, transgenic mice overexpressing PIM3 in the liver showed an increase of lipid droplet accumulation^58^ while PIM1 is known to stabilize the cholesterol transporter^56^ and there is substantial functional redundancy in the PIM kinase family. Accordingly, the high drug concentrations that the liver is exposed to and the unique inhibition of PIM3 by rucaparib (Figure 3) are consistent with the hypothesis that PIM3 kinase inhibition may be responsible for this differential side-effect (Figure 5). However, further experimental and clinical validation is needed to verify this hypothesis, such as comparing biomarkers of PIM3 response in patients treated with different PARP inhibitors.

Interestingly, olaparib does not show off-target activity against any of the kinases tested but its unique side-effects suggest that this drug could have off-target effects beyond kinases. Testing PARP inhibitors side by side in clinical trials and the comprehensive characterisation of their polypharmacology beyond the kinome would be very valuable to further our understanding of their differences at the molecular and clinical levels and to explore opportunities for their maximal exploitation for patient benefit.

The distinct off-target and side-effect profiles between PARP inhibitors that we have identified also caution against the assumption that PARP inhibitors are clinically equivalent in all disease and treatment scenarios. Moreover, any differences could be magnified when PARP inhibitors are used in combination with other drugs that could synergise differently with the different off-target activities of clinical PARP inhibitors. This might be particularly important in drug combinations with immunotherapy due to the likely higher concentrations of PARP inhibitors in the blood compared to the tumour site. These higher concentrations could make the weaker kinase off-target activities capable of impacting the immune response. Currently, there are at least 30 clinical trials studying combinations between PARP inhibitors and immunotherapies (Supplementary Table 10) but none is comparing more than one PARP inhibitor side by side. DYRK1A has been shown to have an important role in immune system homeostasis by regulating the branching point between Th17 and Treg differentiation.^59^ Treg cells are known to modulate the tumour suppressive microenvironment and the prevention of their suppressive activities through immune checkpoint blockade is a major focus of current oncology research.^60^ In this context, the off-target inhibition of DYRK1A may potentially play a role in increasing the Treg cell population that in turn may antagonise the effects in PARP-immunotherapy drug combinations. If this is the case, the combination of olaparib or niraparib with immunotherapy drugs may give different results and we recommend that this should be investigated carefully. Moreover, the submicromolar off-target interactions between rucaparib and PIM3 may also play a role in immune system homeostasis. PIM3 is known to regulate STAT3 by phosphorylating Tyr-705 which is uniquely dephosphorylated after rucaparib treatment at micromolar concentrations in cellular models.^13,61,62^ STAT3 has recently been shown to be involved in the control of responder T-cell senescence induced by human Treg cells.^60^ Consequently, the inhibition of STAT3 phosphorylation by rucaparib via PIM3 may play a role in reducing responder T-cell senescence produced by Treg cells and thus synergise with immunotherapy.^60^ A recent study has also shown that inhibition of PIM kinases improved anti-tumour T cell therapy in an animal model via modulating T cell homeostasis.^63^ However, all the PIM kinases were simultaneously inhibited in the study – and thus these results need to be validated for the selective inhibition of PIM3. Overall, several kinase off-targets of PARP inhibitors could modulate T cell homeostasis and their potential implications should be validated further to maximize PARP drug combinations with immunotherapy.

In summary, our study demonstrates that PARP inhibitors have an inherent capacity to inhibit kinases off-target and illustrates that each of the clinically approved PARP inhibitors investigated in this work has a unique polypharmacological kinase profile. Of particular note, we identify novel submicromolar off-target kinases for rucaparib and niraparib. We demonstrate through our analysis of prescribing information and key clinical trials that FDA-approved PARP drugs have distinct clinical side-effect profiles and we recommend that studies be undertaken to determine the potential contribution of off-target kinase effects to drug side-effects. Our study highlights the field’s currently limited understanding of drug polypharmacology and its implications for efficacy and safety in the clinic. This is particularly important when considering drug combinations with limited understanding of the polypharmacological liabilities of the combined drugs. However, through the application of complementary technologies, we can map key polypharmacological profiles and generate testable hypotheses with clinical potential. In this way, we can help facilitate the maximal exploitation of PARP inhibitors and other drugs for patient benefit.

## ACKNOWLEDGMENTS

This work was primarily supported through the Wellcome Trust Sir Henry Wellcome Postdoctoral Fellowship (204735/Z/16/Z); the People Programme (Marie Curie Actions) of the 7th Framework Programme of the European Union (FP7/2007-2013) under REA grant agreement no. 600388 (TECNIOspring programme) and the Agency of Business Competitiveness of the Government of Catalonia, ACCIO to A.A.A.; B.A.-L., I.C and P.W are funded by The Institute of Cancer Research (ICR) as well as the Cancer Research CRUK programme grant to the Cancer Research UK Cancer Therapeutics Unit (grant C309/A11566). P.W is a CRUK Life Fellow. We acknowledge support from CRUK and the NHS funding to the CRUK Centre and Biomedical Research Centre respectively at the ICR and Royal Marsden NHS Foundation Trust. The authors thank many collaborators for discussions and valuable input into the preparation of this manuscript. In particular, the authors thank Johann de Bono for very valuable discussions and advice.

## METHODS

### In silico target profiling

Three computational methods based on chemical similarity were used to predict the kinase off-targets of clinical PARP inhibitors. The canonical SMILES used to define the chemical structures of the four clinical PARP inhibitors analysed were obtained from ChEMBL.^35^ The first method used was the predefined consensus of six ligand-based chemoinformatic methods available in the Chemotargets CLARITY platform (https://www.chemotargets.com) and a predefined panel including PARPs and kinases was selected for off-target prediction.^34^ Secondly, we used the Similarity Ensemble Approach (SEA) method (http://sea.bkslab.org/) set to default parameters.^26^ And the third method used was the multinomial Naive Bayesian multicategory scikit-learn similarity-based method implemented in ChEMBL that can be accessed from the ChEMBL website (https://www.ebi.ac.uk/chembl/).^35^ The raw data from the predictions we obtained can be accessed in Supplementary Tables 2-4.

### In vitro kinome profiling measuring drug binding

DiscoveRx’s KinomeScan *in vitro* active site-directed competition binding assay (https://www.discoverx.com) was used to quantitatively measure interactions between the four clinical PARP inhibitors and their largest kinase panel. At the time when the assays were performed (date: 29/11/2016), 468 *in vitro* binding assays were available, corresponding to 392 unique human kinases (76% of the human kinome)^39^ (Supplementary Table 5).

### In vitro kinase radiometric assays

Reaction Biology’s HotSpot platform (http://www.reactionbiology.com),^40^ that uses a radioisotope filter binding assay, was used to validate the hits from the kinome binding assay with greater effects at 10 μM. The radiometric assay is designed to directly detect the true product without the use of modified substrates, coupling enzymes, or detection antibodies. Test or control compounds are incubated with kinase, substrate, cofactors, and radioisotope-labeled ATP (32P-γ-ATP or 33P-γ-ATP). The reaction mixtures are then spotted onto filter papers, which bind the radioisotope-labeled catalytic product. Unreacted phosphate is removed via washing of the filter papers.^41^

### Docking experiments

We selected the crystal structures of DYRK1A (PDB ID: 4AZE) and CDK16 (PDB ID: 3MTL) because they were the only ones cocrystalized with a ligand presenting a cyclic benzamide that mirrors the one present in all PARP inhibitors (Supplementary Table 7). The PDB files were prepared using the standard preparation method implemented in GOLD v5.6 (https://www.ccdc.cam.ac.uk/solutions/csd-discovery/components/gold/). The binding site was described selecting residues at a distance of 6 Å from the cocrystalised ligand. The ligand structures were extracted from the PDB and edited to define the correct atom types. Docking was performed using standard variables for high conformation sampling (30 GA runs) and amide bond and ring system flexibility of the ligand were enabled. Best docking poses were analysed with LigPlot+.^44^

### Side-effect analysis

FDA prescribing information was downloaded from the FDA website (www.accessdata.fda.gov; accessed: 24/10/2018). The raw data describing the side-effects and laboratory abnormalities and their frequency was extracted from the FDA prescribing information documents (Supplementary Table 8). A total of 64 side-effects and laboratory abnormalities were described for the four FDA-approved PARP drugs (Supplementary Table 8). From the 64 side-effects, ‘decrease in leucocytes’ and ‘leukopenia’ were considered redundant side-effects and therefore were merged. Similarly, ‘nasopharyngitis / URI / sinusitis / rhinitis / influenza’ and ‘(upper) respiratory tract infection’ were also considered analogous and merged. Finally, ‘ALT increase’ and ‘AST increase’ were also merged in a single side-effect. Accordingly, the final number of side-effects and laboratory abnormalities considered was 61 (Supplementary Table 9). The information was subsequently transformed into a binary format (1 = side-effect present; 0 = side-effect absent) (Supplementary Table 9). Uncommon side-effects by the definition of the FDA label were distinguished from common in two different columns (Supplementary Table 9). Differential side-effects were then compared to larger published clinical trials of olaparib, rucaparib and niraparib.^48^–^50^ Several side-effects that appeared differential only using the FDA labels were found to have been reported in larger clinical trials, such as dyspepsia, headache or myalgia that had been reported for rucaparib in the latest clinical trial despite not being included in the FDA label (Supplementary Table 8-9).^49^ Figure 5 summarises the nineteen side-effects that are different between PARP inhibitors and were also reported as common for at least one PARP inhibitor.

