## Supplementary Figure 1 for "The off-target kinase landscape of clinical PARP inhibitors"

### Supplementary Figures

| Supplementary | Title | Page # |
|---|---|---|
| Supplementary Figure 1 | Docking poses with the top GOLD S(PLC) score of four clinical PARP inhibitors in CDK16 kinase. | 3 |

**Supplementary Figure 1.** Docking poses with the top GOLD S(PLC) score of four clinical PARP inhibitors in CDK16 kinase. In the top left panel, all ligands are superimposed and the whole protein structure is displayed using MacPyMOL (PyMOL v1.8.0.6). In subsequent panels, LigPlot+ was used to generate schematic diagrams of protein-ligand interactions for the cocrystalized ligand and clinical PARP inhibitors. In the bottom, a table summarises the value of the GOLD scoring function for the best pose –that is the one represented– and the IC50 determined in this work (Figure 3) or extracted from the literature for each of the ligands.

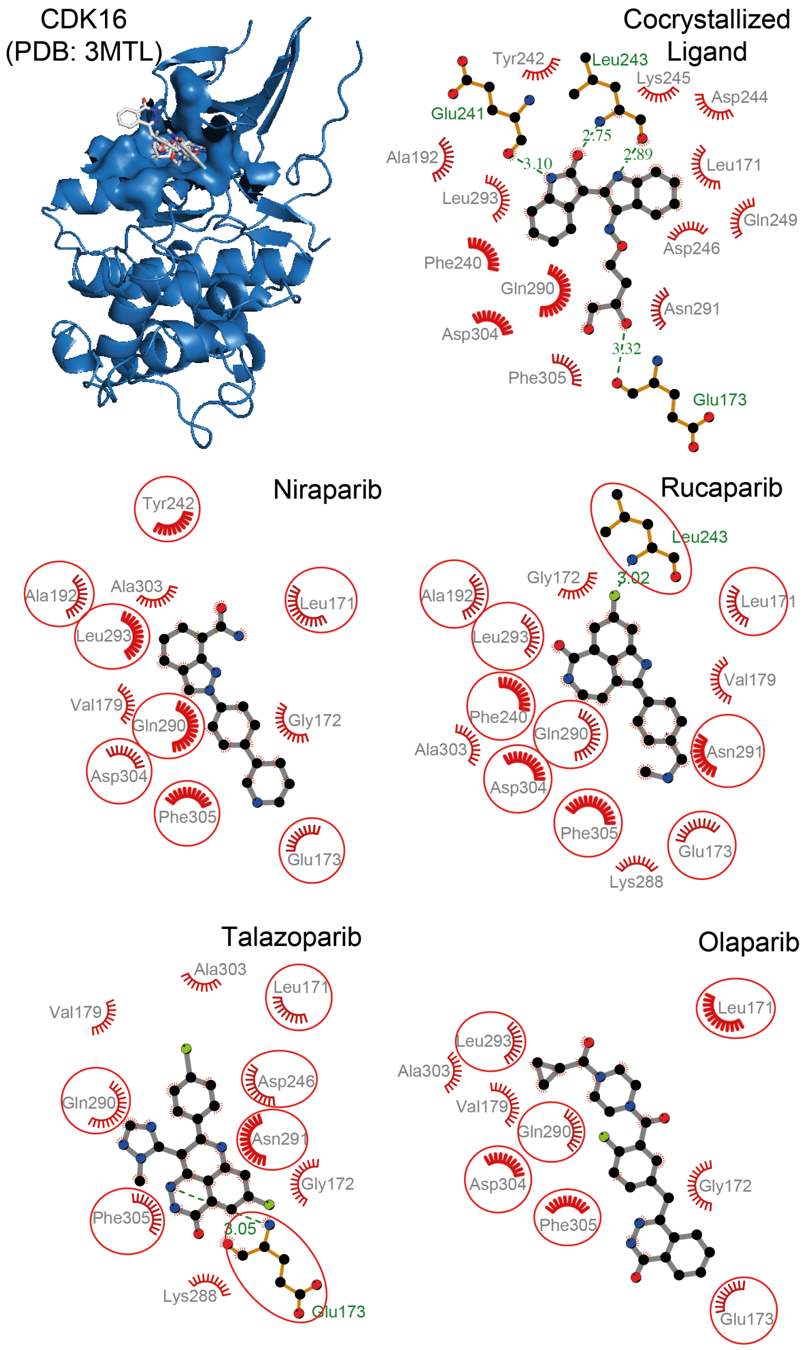


| | Cocrystallized Ligand | Niraparib | Rucaparib | Talazoparib | Olaparib |
|---|---|---|---|---|---|
| IC50 | 83 nM | > 10,000 nM | 381 nM | > 10,000 nM | > 10,000 nM |
| S(PLC) | -80.8 | -58.0 | -60.0 | -42.3 | - 50.4 |
